# SignEEG v1.0 : Multimodal Dataset with Electroencephalography and Hand-written Signature for Biometric Systems

**DOI:** 10.1101/2023.09.09.556960

**Authors:** Ashish Ranjan Mishra, Rakesh Kumar, Vibha Gupta, Sameer Prabhu, Richa Upadhyay, Prakash Chandra Chhipa, Sumit Rakesh, Hamam Mokayed, Debashis Das Chakladar, Kanjar De, Marcus Liwicki, Foteini Simistira Liwicki, Rajkumar Saini

## Abstract

Handwritten signatures in biometric authentication leverage unique individual characteristics for identification, offering high specificity through dynamic and static properties. However, this modality faces significant challenges from sophisticated forgery attempts, underscoring the need for enhanced security measures in common applications. To address forgery in signature-based biometric systems, integrating a forgery-resistant modality, namely, noninvasive electroencephalography (EEG), which captures unique brain activity patterns, can significantly enhance system robustness by leveraging multimodality’s strengths. By combining EEG, a physiological modality, with handwritten signatures, a behavioral modality, our approach capitalizes on the strengths of both, significantly fortifying the robustness of biometric systems through this multimodal integration. In addition, EEG’s resistance to replication offers a high-security level, making it a robust addition to user identification and verification. This study presents a new multimodal *SignEEG v1.0* dataset based on EEG and hand-drawn signatures from 70 subjects. EEG signals and hand-drawn signatures have been collected with *Emotiv Insight* and *Wacom One* sensors, respectively. The multimodal data consists of three paradigms based on mental, & motor imagery, and physical execution: i) thinking of the signaturés image, (ii) drawing the signature mentally, and (iii) drawing a signature physically. Extensive experiments have been conducted to establish a baseline with machine learning classifiers. The results demonstrate that multimodality in biometric systems significantly enhances robustness, achieving high reliability even with limited sample sizes. We release the raw, pre-processed data and easy-to-follow implementation details.

## Background and summary

In biometrics, user *identification* refers to establishing unique credentials or attributes within a system, while user *verification* confirms the claimed identity through interaction with the system. In terms of biometrics, user identification involves establishing the identity of an individual from a larger dataset, while user verification focuses on confirming the claimed identity of an individual through a one-to-one match. Various physical traits, such as retina, iris, sclera, ear, hand, palm geometry, fingerprints, and face, along with behavioral characteristics like keystroke pattern, voice, signature, and gait, have been utilized in the design of biometric systems^1^.

Signature-based biometric systems^2,3^ are preferred due to their convenience, non-invasiveness, cost-effectiveness, wide adoption, cancelable nature, and legal recognition. Biometric authentication^4^ via handwritten signatures leverages the distinct characteristics inherent in an individual’s signature, offering a unique method for identification and verification, particularly in commercial contexts like bank check processing and legal agreements. The intricate process and dynamic motion patterns of each handwritten signature confer a high level of individual specificity, rendering it an effective biometric modality. However, forgery attempts pose a substantial challenge in signature-based biometrics. These forgeries are sophisticated efforts to replicate an individual’s unique signature style, encompassing its dynamic and static characteristics.

To mitigate forgery risks, a viable strategy involves incorporating an additional, user-specific data modality that is inherently resistant to forgery. Simultaneously, this approach would capitalize on the strengths of multimodality, thereby enhancing the overall robustness of the biometric system. Incorporating EEG as a physiological trait and handwritten signatures as a behavioral trait, our approach effectively utilizes the distinct yet complementary aspects of the same individual. This multimodal integration substantially strengthens the robustness of biometric systems, emphasizing the synergy between physiological and behavioral modalities.

EEG-based biometric systems provide a high level of security against forgery due to the inherent difficulty of replicating an individual’s unique brain activity patterns^5^. Unlike signatures or physical biometrics, EEG signals are challenging to forge, making them a reliable and robust method for identity verification.

EEG-based biometric systems^6^ use electroencephalogram (EEG) signals for the automatic recognition of people’s identities. These systems have been successfully used in various states, such as resting, visual stimuli, mental tasks, and emotional stimuli. The extraction of features from EEG signals is a crucial processing step, and a higher quality of identity information is necessary to improve the distinctiveness between subjects in EEG-based biometric systems. EEG-based biometric systems^6^ have several advantages over other neuroimaging techniques. They use relatively simple and inexpensive equipment for signal acquisition and can be used in most environments. The feature extraction and classification techniques of these systems have demonstrated high accuracy rates, even as the number of channels is reduced, and the signal acquisition process becomes less complicated. Given the susceptibility of signature-based biometric systems to sophisticated forgery, integrating EEG capturing unique brainwave patterns impervious to replication would make the biometric system more robust against such fraudulent attempts. There are a few research works focused on integrating the EEG with online^7^ and offline^8^ signatures with confidential data. In addition to the lack of data availability, they investigate EEG in limited settings.

This study integrates brain signals (EEG) and hand-drawn signatures within a multimodal biometric framework to develop SignEEG v1.0 dataset, considering multiple EEG paradigms with different tasks while focusing on two primary objectives that have been defined, namely, *user identification* and *verification*.

The existing research^7,8^ on integrating hand-drawn signatures and EEG signals is scarce and constrained to a single paradigm. Furthermore, the unavailability of publicly accessible data from these studies hinders researchers from further investigations. These EEG and signature-based multimodal works are summarized in Table 1.

**Table 1.** Related Work: EEG and Signatures.

| Study | Method Used | Headset Details | Participants (genuine + forged) | Total Samples | Mental Imagery | Motor Imagery | Physical Execution | Availability |
| --- | --- | --- | --- | --- | --- | --- | --- | --- |
| <sup>7</sup> | wavelet for EEG, PHOG for signs HMM classifier | Emotiv Epoc +,<br>#channel: 14 (+ CMS/DRL)<br>Saline based electrodes<br>Set-up time: 20-30 minutes | 33 + 25 | 1650 | - | - | ✓ | - |
| <sup>8</sup> | DFT for EEG, geometric features for signs BLSTM classifier | Emotiv Epoc +,<br>#channel: 14 (+ CMS/DRL)<br>Saline based electrodes<br>Set-up time: 20-30 minutes | 70 + 70 | 1400 | - | - | ✓ | - |
| SignEEG v1.0 | statistical and wavelet for EEG, statistical for signs traditional machine learning classifiers | Emotiv Insight,<br>#channel: 5 (+ CMD/DRL)<br>Semi-dry polymer sensors with primer fluid<br>Set-up time: 1-2 minutes | 70 + 70 | 2043 × 3 | ✓ | ✓ | ✓ | ✓ |

SignEEG v1.0 consists of three paradigms, unlike others where only one paradigm is used. The proposed dataset gives the opportunity to develop multimodal biometric systems as it covers different paradigms and hand-drawn signatures. By incorporating these three paradigms, a multimodal approach to user authentication can be established, combining the strengths of mental imagery, motor imagery, and physical execution. This comprehensive approach enhances the security and reliability of the biometric system by capturing multiple facets of the individual’s unique signature-related brain activity and behavior. To conclude, our key contributions can be summarized as:

- Our study establishes a novel multimodal approach for developing robust biometric systems by comprehensively encompassing an individual’s traits from both the behavioral aspect, as evidenced by handwritten signatures, and the physiological aspect, as reflected in EEG patterns.
- We develop a multimodal biometric dataset (SignEEG v1.0 ) with EEG and hand-drawn signatures from 70 healthy participants collected over several months.

The rest of the paper is organized as follows. Section *Methods* describes the data acquisition process, including details of participants, tools used, recording setup, experimental paradigms, data acquisition protocol, and objectives, followed by data pre-processing. Sections *Data records, Baseline Results and Discussion*, and *Conclusion* discuss the physical structure of the data, results, and conclusion, respectively.

## Methods

In this section, we describe in detail the data acquisition process, including devices used, participants (subjects), the experimental paradigm, recording setup, and acquisition protocol.

### Participants

The dataset development involved 70 enrolled participants (anonymized). All 70 participants were healthy individuals, comprising five females and sixty-five males, aged between seventeen and thirty-six years, and without any neurological disorders or brain-related medications/surgery. None of the participants had prior experience with EEG. Participants were instructed not to consume alcohol or tobacco for two days prior to the data collection. After the data collection, the participants independently rated their satisfaction, boringness, horribleness, and calmness regarding their experience, and the average scores (out of 10) were 9.56, 1.43, 1.17, and 9.50, respectively. Written informed consent was obtained from all participants before they participated in the experiment; a sample consent form may be provided as supplementary material if required. The ethical approval was obtained from the ethical review board - REC Sonbhadra, U.P., India, with reference number:72*/*13*/REC/SONBH/*2021.

### Device and software details

The **Emotiv Insight headset**^9^ is a wireless EEG device designed for research and personal use. It has five channels *(AF3, AF4, T7, T8, Pz)* for recording EEG signals and two references *(CMS, DRL)*, with a sampling rate of 128 Hz and a resolution of 14 bits. It uses a built-in digital 5^*th*^ order Sinc filter for antialiasing. The electrode placement according to the 10-20 international system^10^ is shown in Figure 1. Colors indicate the quality of the electrical signal passing through the sensors and the reference; colors correspond to the impedance measured by Emotiv’s patented system for both the contact and the signal quality. The impedance ranges between 5-20kΩ, where green and red colored electrodes indicate the lowest and highest impedance values, respectively. The Emotiv Insight headset used Semi-dry polymer material in their sensors; a primer fluid may be used to improve the contact quality with skin^9^.

**Figure 1.**
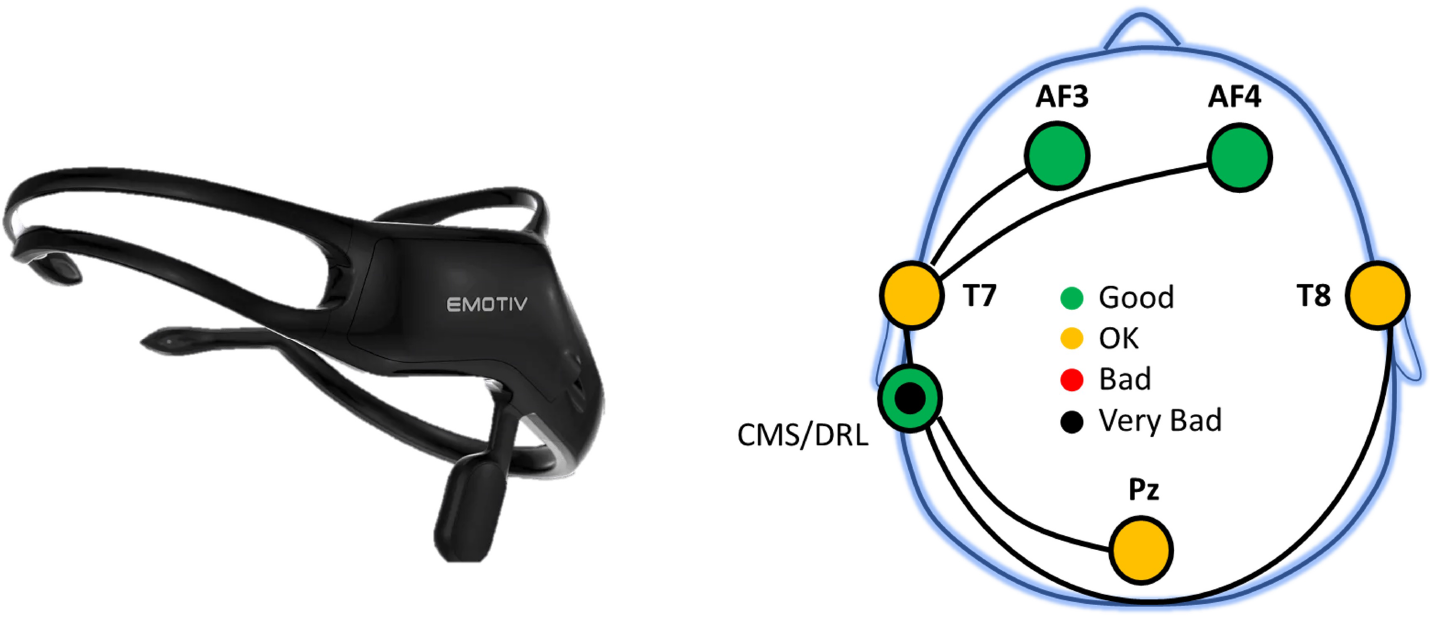
Emotiv Insight headset^9^ (left) and Electrode placement (right) according to the 10-20 international system^10^. The green and orange colors indicate the good and moderate (OK) quality of the signal, respectively. While red and black colors indicate bad and very bad signal quality, respectively. Colors correspond to the impedance measurements. Please refer to the technical specifications for more details^9^.

The headset is lightweight, comfortable, and easy to set up, making it suitable for brain-computer interfaces (BCI) research. The Emotiv Insight requires an **EmotivPRO** software toolkit^11^; it allows researchers to record, visualize, and analyze EEG data collected from the headset. The software provides a user-friendly interface for easy data analysis and visualization. Additionally, the toolkit is quite useful in the development of custom applications and experiments using the Emotiv headset.

The **Wacom One Display Pen Tablet**^12^ has a 13.3-inch Full HD display with a resolution of 1920 *×* 1080 pixels. It supports up to 4096 levels of pressure sensitivity and has a battery-free pen with tilt support. The tablet has a viewing angle of 170 degrees and a color gamut of 72% NTSC (National Television System Committee). It has a durable and scratch-resistant surface and comes with an adjustable stand for comfortable use. The tablet also features HDMI and USB Type-C ports for connectivity. Online signatures drawn on the tablet were captured using the Wacom **SignatureScope** software tool^13^. The software captures real-time signature data and provides trajectory information, including coordinates (x and y), timestamp, stroke, pressure, azimuth, altitude of the pen, and other related information. This information is captured at a sampling rate of 200 frames per second.

### Recording Setup

To ensure a controlled environment for data collection, a physical recording studio was set up, which is illustrated in Figure 2(ii). The studio is designed with acoustic walls to minimize external noise and is maintained at a constant temperature of 20 ^*°*^C. The subjects are seated comfortably and provided with a monitor screen and a Wacom tablet with a stylus for online signature recording. Audio instructions are provided through a speaker to ensure consistency across all subjects. The dataset curator is responsible for monitoring and recording the EEG and online signature data, while an assistant helps the subjects with the EEG headset placement. Both the curator and assistant maintain a safe distance from the subject to avoid any interference with the data collection process. In addition, the studio is thoroughly cleaned and disinfected to ensure hygiene, and the power supply is checked to ensure a stable recording environment before the start of each recording session.

**Figure 2.**
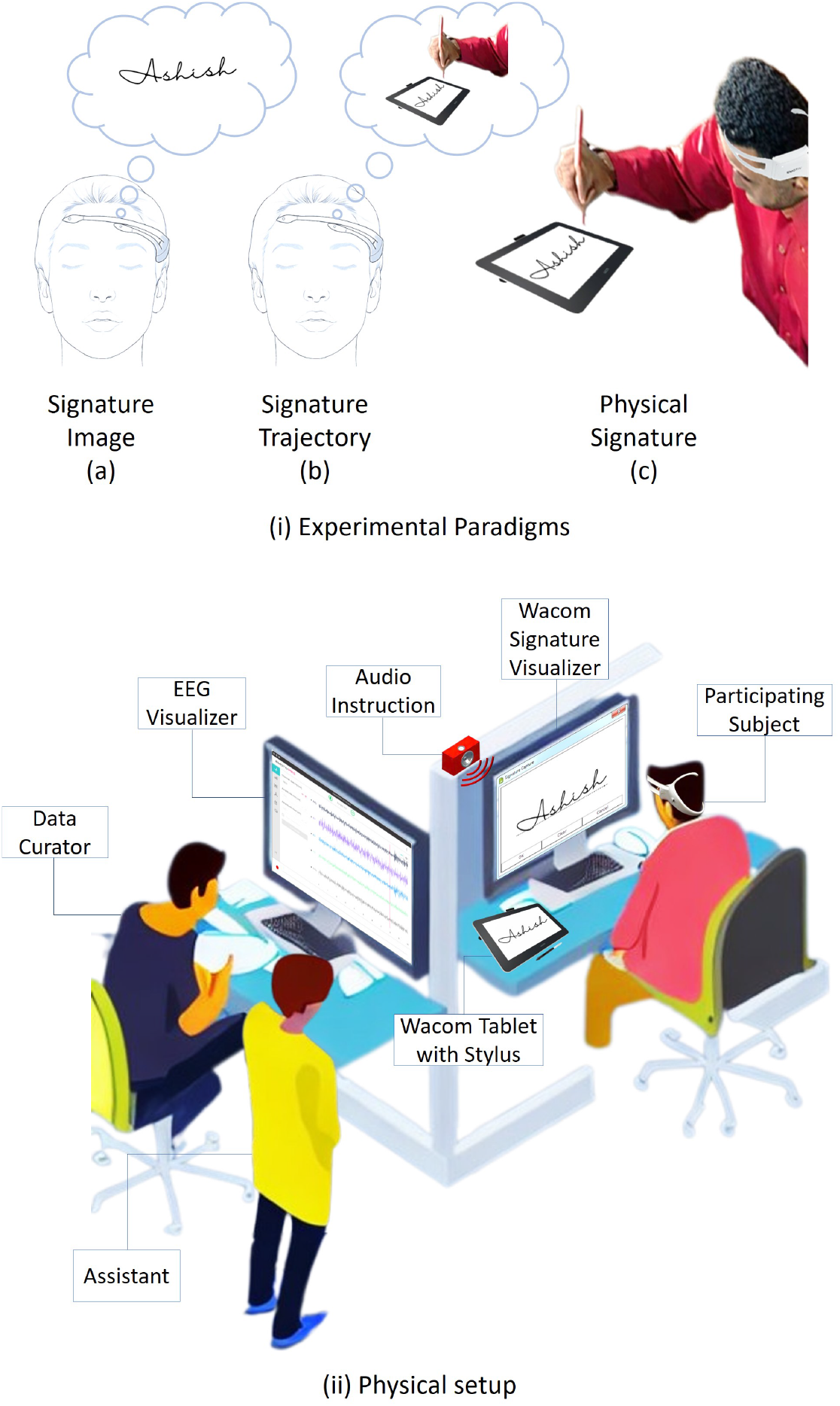
(i) Experimental Paradigms for the multimodal biometric research, (ii) Physical setup of the data collection lab.

### Experimental Paradigms

The multimodal dataset SignEEG v1.0 consists of three paradigms based on mental & motor imagery, and physical execution:

(i) thinking the signaturés image, (ii) drawing the signature mentally, and (iii) drawing the signature physically. These three paradigms are of interest to neuroscience, BCI, and the machine learning research community. During mental & motor imagery in the first two paradigms, participants have their eyes closed while performing the task. In the third paradigm, participants perform the signature on the Wacom tab with open eyes. The acquisition protocol also includes auditory instructions and a rest period before each EEG activity (with eyes closed), as depicted in Figure 3. The three paradigms facilitate researchers in developing unimodal EEG-based biometric systems while having hand-drawn signatures extends the research toward developing multimodal biometric systems. The paradigms are discussed below (also depicted in Figure 2(i)).

**Figure 3.**
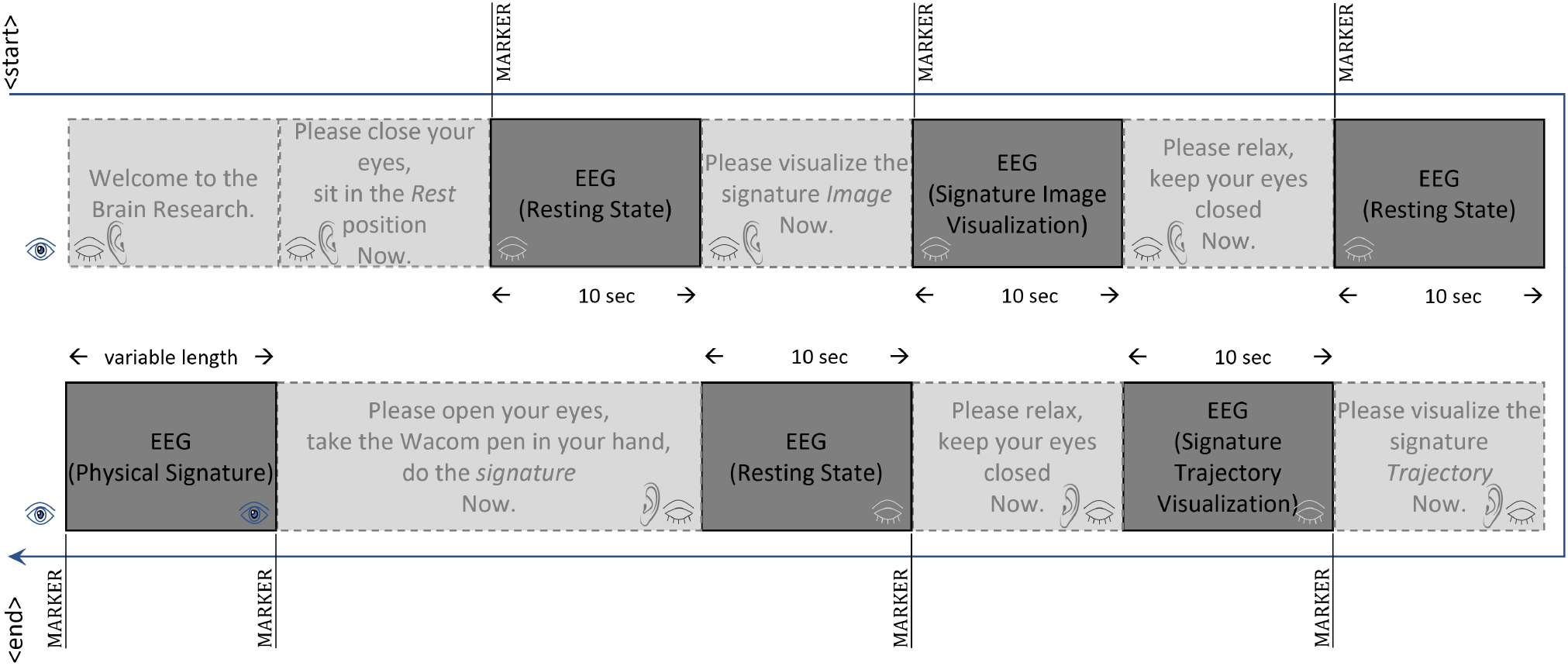
Acquisition protocol for the data collection. Markers (value=4) are inserted into recordings to fetch the EEG signals related to ‘signature’ or ‘rest’ activities.

#### Closed eyes, thinking the signaturés image

This paradigm involves mental imagery^14–16^, the creation of a sensory experience in the absence of external stimuli. When a person closes their eyes and thinks of the signature, they are engaging in mental imagery. This is the simplest of the three paradigms, as it requires minimal motor activity and cognitive effort as compared to others. The main focus is on generating a visual image in the mind’s eye.

#### Closed eyes, drawing the signature mentally

This paradigm also involves mental imagery^14,15^; however, it requires more complex cognitive processes such as spatial reasoning and motor planning. The person needs to create a mental representation of the signature and then simulate the motor movements needed to draw it. This can activate additional brain regions involved in motor planning and execution. This paradigm may also be seen as a motor imagery task from the conventional definition of motor imagery^16^. The detailed discussion can be found in^16,17^.

#### Open eyes, drawing a signature physically

This paradigm involves actual motor execution^18,19^ and visual perception^20^. The person needs to visually perceive the signature and then use motor planning and execution to draw it. This paradigm can activate additional brain regions involved in sensorimotor integration, feedback control, and visual perception. This paradigm requires coordination between the motor and visual systems and engages higher-level cognitive processes such as planning, attention, and working memory.

Different regions of the brain may activate differently in different paradigms; therefore, these paradigms can be analyzed individually or combined.

### Acquisition Protocol

The Emotiv Insight headset^9^ and Wacom One tab^12^ have been used to collect the multimodal SignEEG v1.0 dataset. Wacom tablets (Wacom One and earlier versions) have been used often in handwriting and signature verification research. However, Emotiv Insight is a relatively new headset for BCI research. Researchers have used Emotiv Insight and developed many applications. The study^21^ explored emotion recognition using EEG signals, focusing on the identification of happy and sad states through the analysis of brain signal features and power spectral density derived from two EEG channels. The research utilized data from 26 subjects, applying preprocessing and Empirical Mode Decomposition, followed by classification via an artificial neural network with an accuracy of 88.5%. Similarly, Olivares-Figueroa et al.^22^ proposed a BCI method to monitor emotional states in drone pilots, using EEG data and a database classifying ‘Quiet’ and ‘Very Tense’ states with an algorithm for real-time artifact removal and feature extraction for classification through K-Nearest Neighbors (KNN) and Support Vector Machine (SVM), achieving notable accuracy in emotional state differentiation. Zabcikova^23^ evaluated the Emotiv Insight headset for measuring brain responses to visual and auditory stimuli, utilizing the Emotiv Xavier ControlPanel software. Results indicated significant variations in engagement, stress, and excitement levels among participants responding to different stimuli. The findings indicated that the Emotiv Insight device is well-suited for entertainment applications.

The acquisition protocol is designed based on mental & motor imagery and physical execution of the activity performed by the subjects. This would allow researchers to develop EEG-based unimodal systems and multimodal systems with signatures included. An acquisition protocol has been established to facilitate the SignEEG v1.0 multimodal dataset collection. The protocol employed for recording both EEG and online signature data is illustrated in Figure 3. Initially, subjects sit in a calm/resting state; the subject’s head is fitted with the EEG headset, and electrodes are placed on the scalp according to the 10-20 system^10^. After the channel and EEG signal quality indicators reach a stable state (indicated by the green color in the Emotiv Pro software), the EEG data acquisition is initiated following the procedure outlined in Figure 3.

During the EEG data collection, except for the period when the subjects are providing their physical signatures, their eyes are closed. The subjects are instructed via audio commands throughout the data collection process, with the audio instructions playing simultaneously with the EEG recording. At the start of the data collection, the audio instructs the subjects to ‘Welcome to the Brain Research. Please close your eyes, sit in the Rest position Now’. Following this audio instruction, a marker with a value of 4 is inserted into the EEG recording, signifying the beginning of the rest period. During this rest period, which lasts for 10 seconds, the subjects are instructed to maintain a resting position with their eyes closed.

Subsequently, the audio command instructs the subjects to ‘Please visualize the signature Image Now,’ and a second marker is inserted into the EEG recording. During the following 10 seconds, the subjects are instructed to visualize the image of their signature in their minds’ eye, and if the image disappears, they are to recall it in their minds again. After the 10 seconds have elapsed, the audio instruction advises the subjects to ‘Please relax, keep your eyes closed Now.’ Following this instruction, a third marker is inserted into the EEG recording, and the subjects are instructed to remain in a resting position for an additional 10 seconds.

Subsequently, the audio instruction directs the subjects to ‘Please visualize the signature Trajectory Now.’ For the signature trajectory task, the subjects are instructed beforehand to simulate the act of signing on paper in their minds. They are required to envision the process of producing the signature from start to finish as if they were signing on a physical sheet of paper. Following this instruction, a fourth marker is inserted into the EEG recording, and the subjects simulate the signature as many times as they can within the next 10 seconds. After the 10 seconds have passed, the audio command advises the subjects to ‘Please relax, keep your eyes closed Now.’ Following this instruction, a fifth marker is inserted into the EEG recording, and the subjects are required to remain in a resting position for a further 10 seconds.

Following the resting period, the audio instruction prompts the subjects to ‘Please open your eyes, take the Wacom pen in your hand, do the signature Now.’ After the audio instruction, the subjects opened their eyes and picked up the Wacom stylus in their hand. Once they are ready to perform the signature on the Wacom tablet, a sixth marker is inserted into the EEG recording. The subjects proceed to sign on the Wacom tablet, and the final marker (7th time) is inserted into the EEG recording, indicating that the signature process has been completed. At this point, the dataset curator saves both the EEG and online signature data into permanent storage.

All markers inserted into the EEG recordings, except for the last one (7th marker), indicate the beginning of the EEG activities, whether it is the resting period or related to the signature tasks. It is noteworthy that all markers in the EEG recordings are represented by the digit 4. Additionally, in an EEG recording, each of the EEG activities, except for the one corresponding to the physical signature task, has a duration of 10 seconds. The final EEG activity associated with the physical signature has a duration equivalent to the time spent performing the signature on the Wacom tablet.

### Objectives and Tasks

The complete data acquisition process intends to explore two major objectives: user identification and verification. *User Identification* involves establishing the identity of an individual from a larger dataset. To identify who the subject is from the pool, given the input data. *User Verification* focuses on confirming the claimed identity of an individual through a one-to-one match. Given input data, it aims to verify if the subject is *Genuine* or *Forged*. The following tasks are designed to evaluate the above-mentioned objectives for different experimental paradigms :

*Task 1*: This task aims to develop a biometric system based on paradigm one that corresponds to the EEG signals of mental imagery of the participant’s signatures.

*Task 2*: This task aims to develop a biometric system based on paradigm two that corresponds to the EEG signals of motor imagery of the participant’s signatures.

*Task 3*: This task aims to develop a biometric system based on paradigm three that corresponds to the EEG signals of the physical execution of the signatures.

*Task 4*: This task aims to develop a biometric system combining paradigm three and the Wacom signatures.

These tasks encourage the researchers to develop unimodal and multimodal biometric systems with EEG signals and Wacom signatures.

### Data Pre-processing

The pre-processing of the EEG data was done with EEGLAB^24^ (a MATLAB-based toolbox) following the EEG data pre-processing pipeline as depicted in Figure 4. Epoching refers to segmenting the continuous EEG data into smaller segments for further analysis. The EEG data obtained using the acquisition protocol underwent further processing before being subjected to signal processing. For each subject, there were instances of genuine and forgery attempts (approximately 30 in total) recorded. EEG activities corresponding to rest and signatures were extracted by identifying inserted markers (marker value = 4). Each of the EEG activities related to the first six markers had a duration of 10 seconds, equivalent to 1280 frames starting from the marker’s position, given that the EEG headset had a sampling rate of 128 frames per second. The duration of the last EEG activity varied depending on when the subject completed the signature on the Wacom tablet.

**Figure 4.**
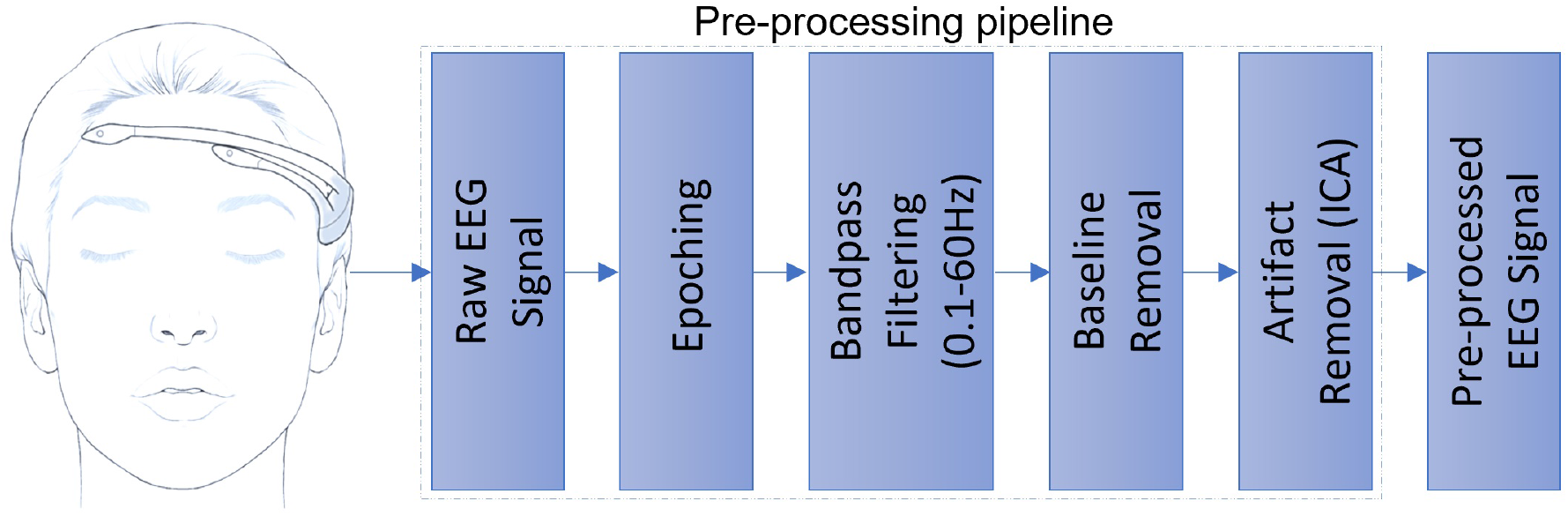
EEG data pre-processing pipeline.

## Data records

The data are available in Zenodo^25^. In total, 2’047 (*×*3, *edf, csv, mat*) instances have been recorded and pre-processed. Each subject is given a unique number (e.g., *200894*). The directory structure ‘*200894\ Genuine*’ and ‘*200894\ Forged*’ consists of instances of genuine and forgery attempts for the subject *200894*, respectively. The file naming convention is as follows. **EDF**: SubjectID_{‘*Genuine\Forged*’}_SignerID_*d*.edf. This is a recorded EEG datum in standard EDF format. The digit *d* represents the instance number. A signer (*SignerID*) performs the EEG and signatures for the subject (*SubjectID*). *SubjectID* and *SignerID* are the same for a *Genuine* attempt, and they differ for a forgery (*Forged*) attempt. The EDF file contains the EEG signals corresponding to the channels in the Emotiv Insight headset, along with other EEG-related information. **CSV**: SubjectID_ ‘*Genuine\Forged*’}_SignerID_*d*.csv. This is a recorded Wacom signature datum in standard CSV (comma-separated value) format. It follows the same naming convention as EDF except for the extension. It consists of the pen data (Index, Stroke, Btn, coordinate X, coordinate Y, Timestamp (T), Pressure, Azimuth, Altitude) per frame.

**MAT**: SubjectID_{‘*Genuine\Forged*’}_SignerID_*d*_epochLength_diff.mat. This is a MATLAB structure array (unordered *struct*, with MATLAB variable name *subject*) consisting of data related to EEG and Wacom signature after pre-processing. It follows the same naming convention as EDF; however, it has additional information in the filename. *epochLength* is the epoch length, and *diff* is an averaged absolute difference of EEG signals before and after applying ICA. The fields of the MATLAB structure and their description are given in Table 2.

**Table 2.**
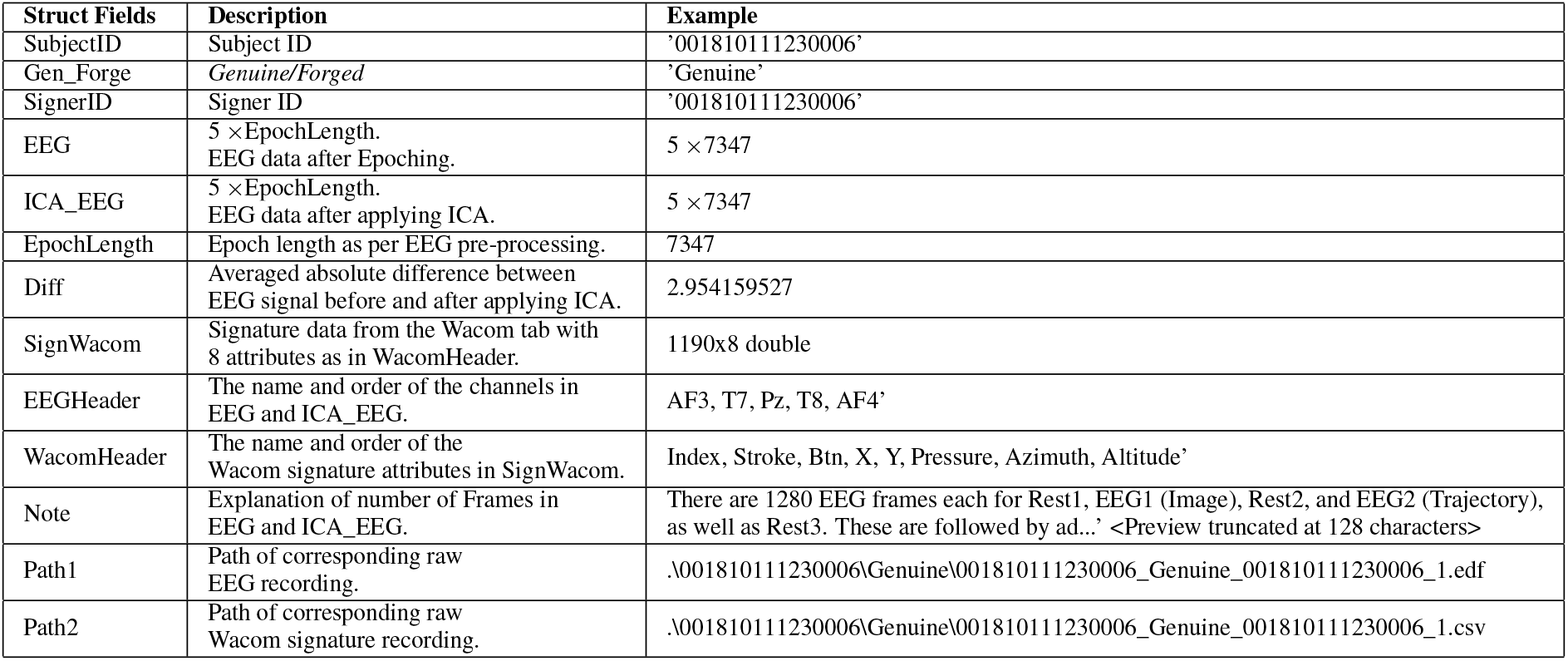
MATLAB structure of pre-processed data.

| Struct Fields | Description | Example |
| --- | --- | --- |
| SubjectID | Subject ID | '001810111230006' |
| Gen_Forge | *Genuine/Forged* | 'Genuine' |
| SignerID | Signer ID | '001810111230006' |
| EEG | $5 \times \text{EpochLength}$.<br>EEG data after Epoching. | $5 \times 7347$ |
| ICA_EEG | $5 \times \text{EpochLength}$.<br>EEG data after applying ICA. | $5 \times 7347$ |
| EpochLength | Epoch length as per EEG pre-processing. | 7347 |
| Diff | Averaged absolute difference between<br>EEG signal before and after applying ICA. | 2.954159527 |
| SignWacom | Signature data from the Wacom tab with<br>8 attributes as in WacomHeader. | 1190x8 double |
| EEGHeader | The name and order of the channels in<br>EEG and ICA_EEG. | AF3, T7, Pz, T8, AF4' |
| WacomHeader | The name and order of the<br>Wacom signature attributes in SignWacom. | Index, Stroke, Btn, X, Y, Pressure, Azimuth, Altitude' |
| Note | Explanation of number of Frames in<br>EEG and ICA_EEG. | There are 1280 EEG frames each for Rest1, EEG1 (Image), Rest2, and EEG2 (Trajectory),<br>as well as Rest3. These are followed by ad...' <Preview truncated at 128 characters> |
| Path1 | Path of corresponding raw<br>EEG recording. | .001810111230006\Genuine\001810111230006_Genuine_001810111230006_1.edf |
| Path2 | Path of corresponding raw<br>Wacom signature recording. | .001810111230006\Genuine\001810111230006_Genuine_001810111230006_1.csv |

Out of 70, the data of 5 subjects is kept aside for robustness analysis in anticipated future research.

## Technical Validation

It is recommended that the epoch length be consistent for a specific type of EEG activity. Therefore, the duration of the last EEG activity, which corresponds to physical signatures, is constant in all instances (genuine and forged) for a given subject.

The instances corresponding to the last EEG activity for each subject were interpolated^26^ to match the length of the longest genuine instance for that subject. This means that all instances (genuine and forged) for a particular subject have the same length (epoch length = 5 *×* 10 *×* 128 +*k*), where *k* is the duration of the last EEG activity. The value of *k* varies among subjects. Once the EEG data has been segmented using epoching, a bandpass filter is applied to the data, with a frequency range of 0.1-60 Hz, followed by the baseline removal. The purpose of baseline removal is to eliminate non-neural activity in the EEG signal by subtracting the mean of the signal from its data points. This process results in a more refined signal that reflects the neural activity under investigation. Following epoching and filtering, Independent Component Analysis (ICA) is utilized to remove artifacts resulting from various sources such as eye movements, muscle activities, and environmental noise.

Toa et al.^27^ and Heunis^28^ have also used ICA on Emotiv Insight data to remove the noise from the EEG signals. By applying ICA, the components associated with the artifacts, as well as the brain activity, are identified. The components with a substantial amount of artifacts are eliminated, while the components linked with the brain activity are kept to produce the final pre-processed signals. The brain maps (corresponding to components (IC) 2, 3, and 4) for two subjects are depicted in Figure 5. It can be noticed from the figure that substantial brain data is preserved while the artifacts are substantially eliminated. A signal before (black) and after (red) the application of ICA is shown in Figure 6, indicating that artifacts are substantially reduced. Besides traditional ICA-based methods, deep learning-based approaches^29,30^ can also be used for artifact removal from EEG signals.

**Figure 5.**
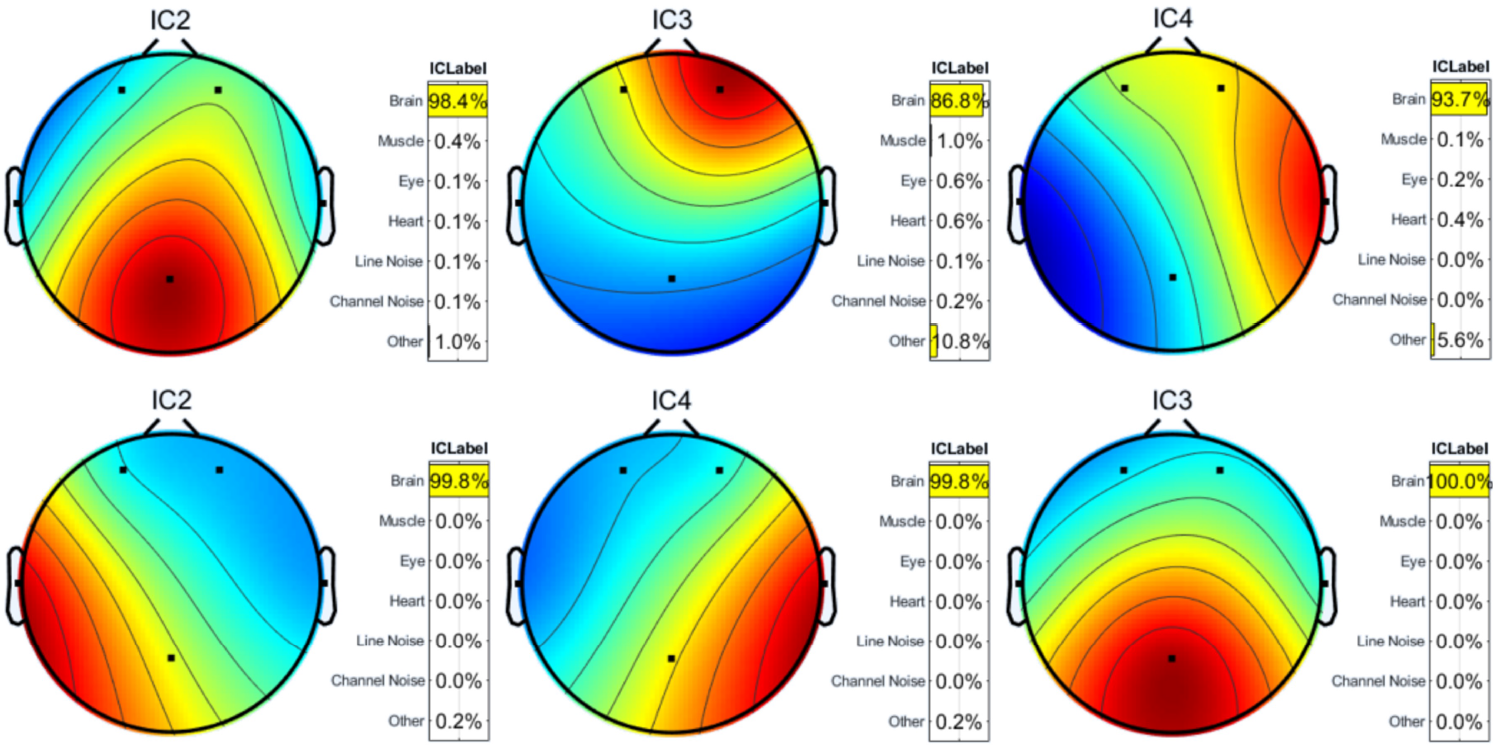
Brain maps corresponding to IC 2, 3, and 4 for two subjects (row-wise). Red color indicates high brain activity.

**Figure 6.**
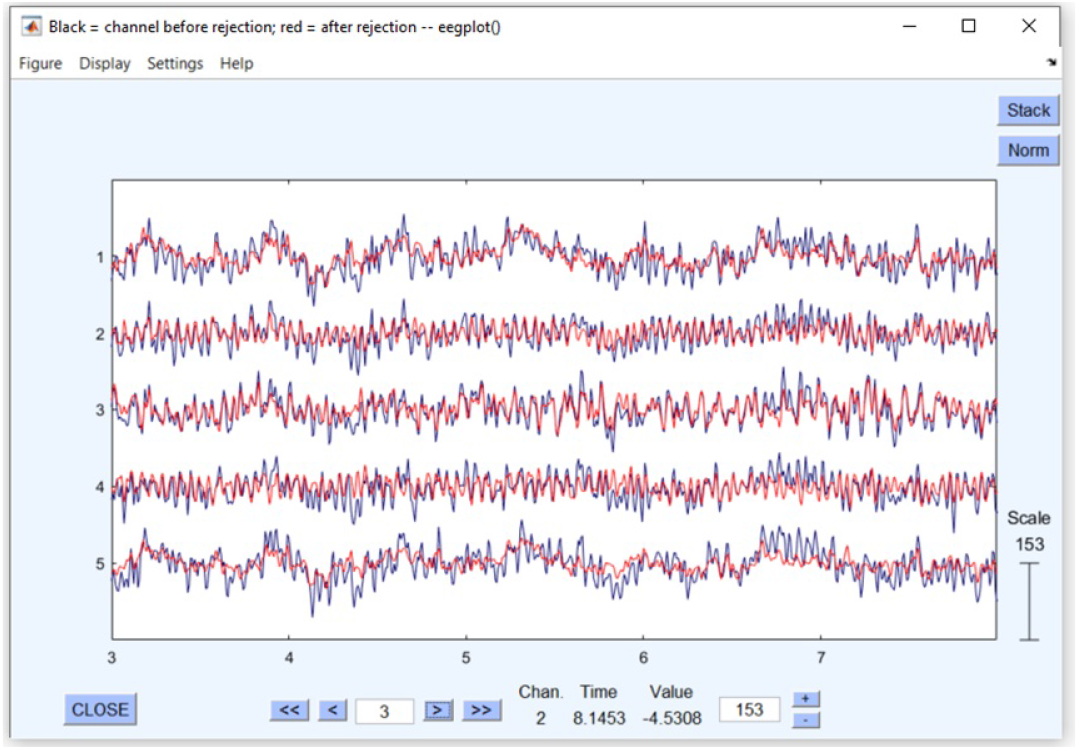
A signal before (black) and after (red) applying ICA in EEGLAB.

### Baseline Results and Discussion

#### Models and features

Table 3 summarizes the performance of several models on two independent objectives, namely identification and verification.

**Table 3.**
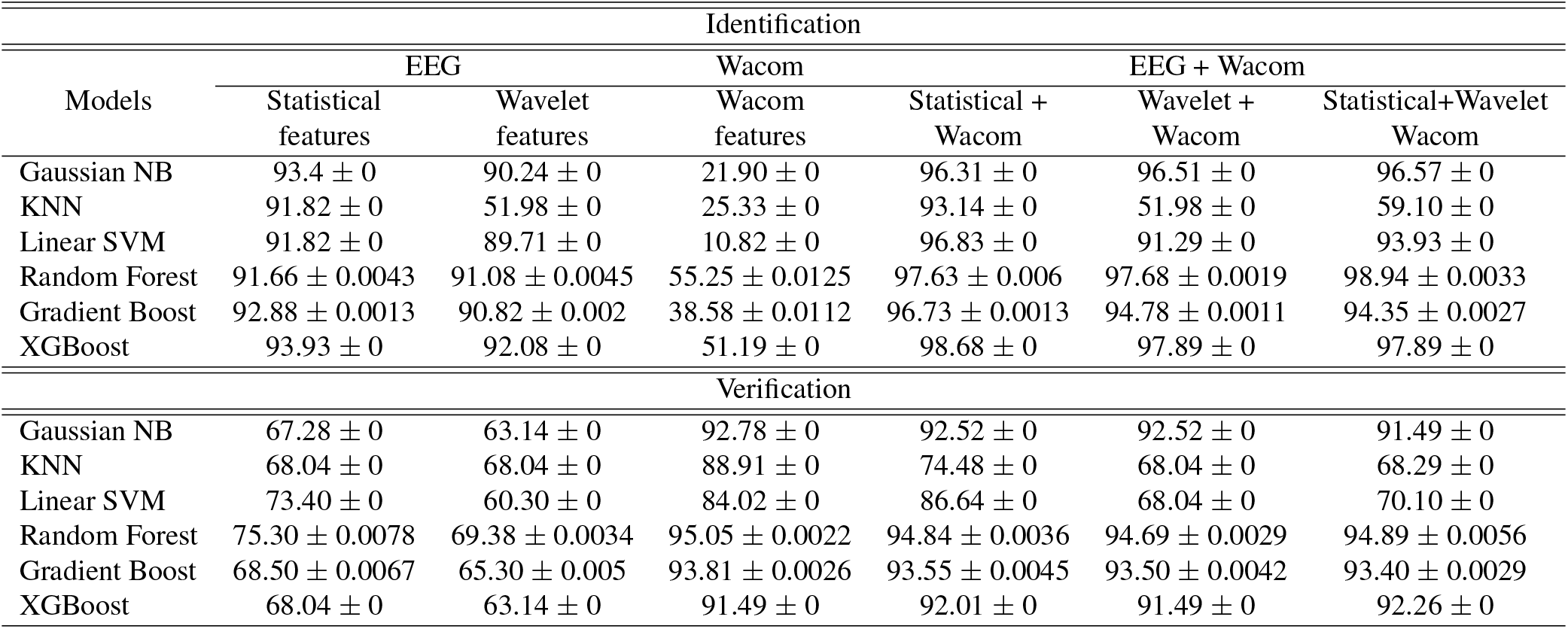
Baseline results on various machine learning and deep learning models for identification and verification. The accuracy score in mean and standard deviation for five trials of each experiment are shown. This is a performance analysis on the data of Task-4, i.e., EEG collected while physically drawing the signature on the Wacom tablet.

For verification, Table 3 represents the combined accuracy across all the subjects, i.e., combining all the true positives, true negatives, and so on. The models include Gaussian Naive Bayes (NB), K-Nearest Neighbors (KNN), Linear Support Vector Machine (SVM), Random Forest, Gradient Boost, and XGBoost. These models were evaluated using different sets of features derived from electroencephalogram (EEG) data (statistical and wavelet)^31^, Wacom tablet data^32^, or a combination of both. Feature extraction is a process in which relevant information is extracted from raw data to represent it in a more compact and meaningful form. Feature extraction refers to the process of identifying and quantifying distinctive patterns or characteristics that are informative for both identification and verification objectives.

The feature extraction process from EEG signals involves normalizing (Z-score normalization) the data channel-wise to ensure consistency. Several statistical features^31^ are derived from the normalized data, including the sum of values, energy, standard deviation, root mean square (RMS), skewness, and kurtosis. These features capture the overall activity, power, variability, magnitude, asymmetry, and peakedness of the signals. Wavelets have also been used to extract wavelet-based features^31,33^ from EEG signals. In this work, the DB4 wavelet is used for the wavelet decomposition of the EEG signals.

The resulting wavelet coefficients are normalized, and the same statistical features are extracted from these coefficients. This approach captures both time-domain and frequency-domain characteristics, providing a comprehensive set of features that enhance the analysis and classification of EEG data.

The feature extraction process from Wacom signature data involves calculating various statistical and dynamic measures^34,35^ to comprehensively capture the signature’s characteristics. Firstly, the raw data, including X and Y coordinates, pressure, azimuth, and altitude, is normalized (Z-score normalization) to standardize the measurements across different sessions. From the normalized data, the first derivatives, representing speed, and the second derivatives, representing acceleration, are calculated for the X and Y coordinates. This allows for the assessment of the movement dynamics. Several statistical features are then extracted from these measures. Specifically, for each of the five parameters—speed, acceleration, pressure, azimuth, and altitude—the mean, standard deviation, and maximum value are computed. The mean provides an average value, reflecting the general level of the measure. The standard deviation indicates the variability or dispersion, showing how much the values fluctuate. The maximum value captures the peak level attained, indicating the most extreme observed value. Additionally, specific features such as the maximum speed and maximum acceleration are explicitly calculated to highlight the highest rates of change in position and velocity. Experiments on EEG and Wacom data have been conducted in different feature settings, as stated in the Table 3 header.

#### Results and discussion

The data has been split into train (60%), validation (20%), and test (20%) sets for the experiments. For the statistical features derived from EEG signals in time domain, the Gaussian (NB), (KNN), Linear SVM, and random forest models exhibit similar performance with accuracy in the low 90% range. Gradient boost and XGBoost also perform well, with accuracies of 92.88% and 93.93%, respectively. Among these, XGBoost performs the best. While on the wavelet features, the performance of KNN is unsatisfactory. Of all the rest of the models, gradient boost and XGBoost perform relatively better, with an accuracy of around 90%. Compared to EEG signal-based features, the model performs less optimally for features from the Wacom signal. Random forest gives the highest accuracy at 55.25%. The combination of statistical and Wacom features drastically improves the performance of all models. XGBoost shows the highest accuracy at 98.68%, followed closely by Random Forest at 97.63%. All other models also exhibit significant improvement in their accuracies. Similar improvements in performance can be observed when wavelet and Wacom features are taken into account. However, KNN does not seem to benefit significantly from this combination, maintaining a low accuracy of 51.98%. The combination of EEG statistical, EEG wavelet, and Wacom features gives most models the highest accuracy, with random forest delivering the best results (98.94%). The only exception is KNN, which only achieves 59.10% accuracy.

In general, it has been noted that the accuracy of KNN is suboptimal compared to alternative models. This may be attributed to the selection of the parameter *K*, which denotes the number of neighbors to be taken into account during the prediction process. In order to ensure fair evaluation, a consistent value of *K* (i.e., 5) has been utilized across all features. However, this may prove inadequate for wavelet and Wacom features, resulting in a decline in performance. On the other hand, XGBoost and random forest excel for most of the experiments, whether statistical or wavelet features. XGBoost offers several advantages that may explain its superior performance in handling various types of features, such as in-built L1 and L2 regularization and cross-validation, tree pruning, capability to handle missing as well as sparse data, and many more. The random forest ensemble learning method shows a strong performance due to its ability to mitigate overfitting through the aggregation of results from multiple decision trees. The pattern in the performance across different models and features seen during identification also appears during verification. Random forest and gradient boost both achieve the highest accuracy when using statistical, wavelet, and Wacom features combined, with accuracies of 94.89% and 93.40%, respectively.

This paper focuses on multimodal biometrics with noninvasive EEG-based brain signals and hand-drawn signatures. However, the medical community has used invasive methods for brain-to-text communication. In^36^, authors used an intracortical Brain-Computer Interface (BCI) that decodes attempted handwriting movements from neural activity in the motor cortex and translates it to text in real-time, using a recurrent neural network decoding approach. They have achieved typing speeds of 90 characters per minute with 94.1% accuracy for individuals whose hand was paralyzed from spinal cord injury.

Besides, noninvasive methods with EEG have also been developed in robotic research. In^37^, authors developed a noninvasive approach using EEG to control a robotic device, improving user engagement and the quality of neural data, which greatly enhances learning and control in realistic tasks. They have achieved nearly 60% accuracy with basic tasks and over 500% in continuous pursuit tasks and could significantly advance the use of BCIs in everyday settings.

### Limitation

In this study, we propose the SignEEG v1.0 dataset combining behavioral (handwritten signatures) with physiological (EEG) data to develop robust biometric identification and verification systems. Alongside traditional unimodal tasks, the SignEEG v1.0 dataset uniquely includes a task dedicated to the development of a multimodal biometric system. The results from our extensive empirical evaluation conclusively demonstrate that the integration of signatures and EEG in a multimodal framework significantly enhances system robustness, effectively utilizing the combined strengths of both modalities. Our study reveals that the combination of behavioral biometrics, exemplified by handwritten signatures, with physiological metrics, represented by EEG, confers a significant advantage to our multimodal approach. The interplay between these two distinct modalities not only complements but also reinforces each other, underscoring the efficacy of this integrated system.

The longitudinal temporal dynamics of EEG are yet to be explored. Therefore, in the future, we will intensively explore the temporal dynamics of EEG and handwritten signatures in a multimodal context through prolonged, longitudinal data collection from the same subjects. Emphasizing the temporal aspect, this approach aims to reveal how these biometric modalities evolve and interact over time, offering critical insights for advancing multimodal biometric system accuracy and reliability.

## Code Availability

The data and source code used in this study are distributed under the Creative Commons Attribution 4.0 International (CC BY 4.0) License (https://creativecommons.org/licenses/by/4.0/) and are available at Zenodo^25^.

